# Augmented base pairing networks encode RNA-small molecule binding preferences

**DOI:** 10.1101/701326

**Authors:** Carlos Oliver, Vincent Mallet, Roman Sarrazin Gendron, Vladimir Reinharz, William L. Hamilton, Nicolas Moitessier, Jérôme Waldispühl

## Abstract

**Motivation:** The binding of small molecules to RNAs is an important mechanism which can stabilize 3D structures or activate key molecular functions. To date, computational and experimental efforts toward small molecule binding prediction have primarily focused on protein targets. Considering that a very large portion of the genome is transcribed into non-coding RNAs but only few regions are translated into proteins, successful annotations of RNA elements targeted by small-molecule would likely uncover a vast repertoire of biological pathways and possibly lead to new therapeutic avenues.

**Results:** Our work is a first attempt at bringing machine learning approaches to the problem of RNA drug discovery. `RNAmigos` takes advantage of the unique structural properties of RNA to predict small molecule ligands for unseen binding sites. A key feature of our model is an efficient representation of binding sites as augmented base pairing networks (ABPNs) aimed at encoding important structural patterns. We subject our ligand predictions to two virtual screen settings and show that we are able to rank the known ligand on average in the 73rd percentile, showing a significant improvement over several baselines. Furthermore, we observe that graphs which are augmented with non-Watson Crick (a.k.a non-canonical) base pairs are the only representation which is able to retrieve a significant signal, suggesting that non-canonical interactions are an necessary source of binding specificity in RNAs. We also find that an auxiliary graph representation task significantly boosts performance by providing efficient structural embeddings to the low data setting of ligand prediction. `RNAmigos` shows that RNA binding data contains structural patterns with potential for drug discovery, and provides methodological insights which can be applied to other structure-function learning tasks.

**Availability:** Code and data is freely available at http://csb.cs.mcgill.ca/RNAmigos.

## 1 Introduction

Recent studies have identified small organic molecules as important non-covalent regulators of RNA function in many cellular pathways [1]. These discoveries contribute to a better understanding of key pathways present in all organisms, but also pose RNA molecules as a large class of promising novel drug targets. For example, Ribocil, which has recently been uncovered through a phenotypic assay to target the FMN riboswitch, is currently undergoing clinical trials as a novel antibiotic [2]. Various other small molecule-activated RNA systems are also being proposed [3, 4, 5]. Notable among these is the application to CRISPR activation regulation [6]. The list of possible therapies is likely to expand given the observations of KD Warner and co-workers that only a small fraction of the genome is translated into protein (1.5%) while the vast majority is transcribed into potentially druggable non-coding RNA (70%) [7].

In parallel, the protein-binding drug discovery field is experiencing rapid and significant advances from data-driven artificial intelligence models. Tasks such as candidate molecule generation and affinity scoring [8] which were previously done using costly knowledge-based simulations are now highly automated [9]. One of the largest points of resistance for data-driven RNA drug discovery is the relatively small number of available binding assays and crystallized RNA-small molecule complexes (34,271 solved protein structures with a ligand, vs 2,253 for RNA). While the number of solved RNA structures is steadily growing, successful models for automated RNA 3D function annotation will likely require customized methods which carefully leverage domain knowledge. To this end, we turn to current knowledge of RNA structural organization to motivate base pairing networks as a strong prior in low-data settings.

### 1.1 RNA Structural Organization

RNAs possess multiple levels of structural organization which together determine function, and by extension, ligand binding specificity. At the simplest level, RNA is a string of monomers {`A,U,C,G`} linked by a chain of covalent bonds known as the backbone. This is commonly known as as the primary structure of RNA. Non-covalent pairwise interactions between nucleotides (bases) in the chain give rise to the secondary and tertiary structure. Canonical pairs (i.e. `A-U, C-G`), are the most studied class of base pair, give rise to the secondary structure. Notably, canonical pairs series of nested loops and stable stacks, assembling a scaffold for the full structure [10]. The experimental determination of binding energies for these pairs [11] prompted a boom of explicit algorithms for sequence to secondary structure prediction such as RNAfold, [12]. In seminal work, Leontis and Westhof identified 11 additional types of base pairing occurring in 3D structures [13, 14], known as non-canonical base pairs. These interactions can occur between any pair of nucleotides and are defined by the relative orientations of three faces of the interacting bases. By considering all combinations of faces and a *cis* and *trans* orientation, we arrive at 12 possible base pairing geometries. Whereas canonical pairs form stable helices, non-canonical pairs are enriched in loops (i.e. regions without canonical pairs) and create more complex structural patterns [15, 16]. These pairings fine-tune RNA function by defining structure at the 3D level [17]. Interestingly, non-canonical pairs were also found to be enriched in ligand binding sites [18, 19], which corroborates with the observation that increased structural complexity is associated with binding specificity [7].

These observations led us to hypothesize that studying RNA structures at the augmented base-pairing level (i.e. including non-canonical pairs) holds useful spatial and chemical information about ligand binding. However, studying RNA at this level of structure comes with major algorithmic challenges, namely, binding energies of non-canonical interactions has not yet been determined, and non-canonical interactions are known to be highly non-nested (i.e. many crossing interactions). For these reasons, non-canonical interactions are typically modeled with statistical methods, and represented using more general data structures such as graphs [20] instead of trees or grammars as is the case for 2D structures. In practice, this means that a graph using vertices to represent nucleotides and multi-relational edges to encode base-pairing interactions could offer a signature for RNA ligand binding sites (See **Fig. 1** for an example of a binding site and its associated base pairing network). We call this graphical representation of RNA sites annotated with canonical and non-canonical interactions an Augmented Base Pairing Network (ABPN) since we consider base pairs beyond the canonicals. Indeed, similar representations of RNA base pairing networks have been exploited in various tools [21, 22, 16, 20] for their ability to capture RNA-specific interactions in an interpretable manner. This paradigm distinguishes RNA from protein-ligand interactions where surface-cavity topologies tend to drive binding preferences [23], making direct use of atomic coordinates more appropriate.

**Figure 1:**
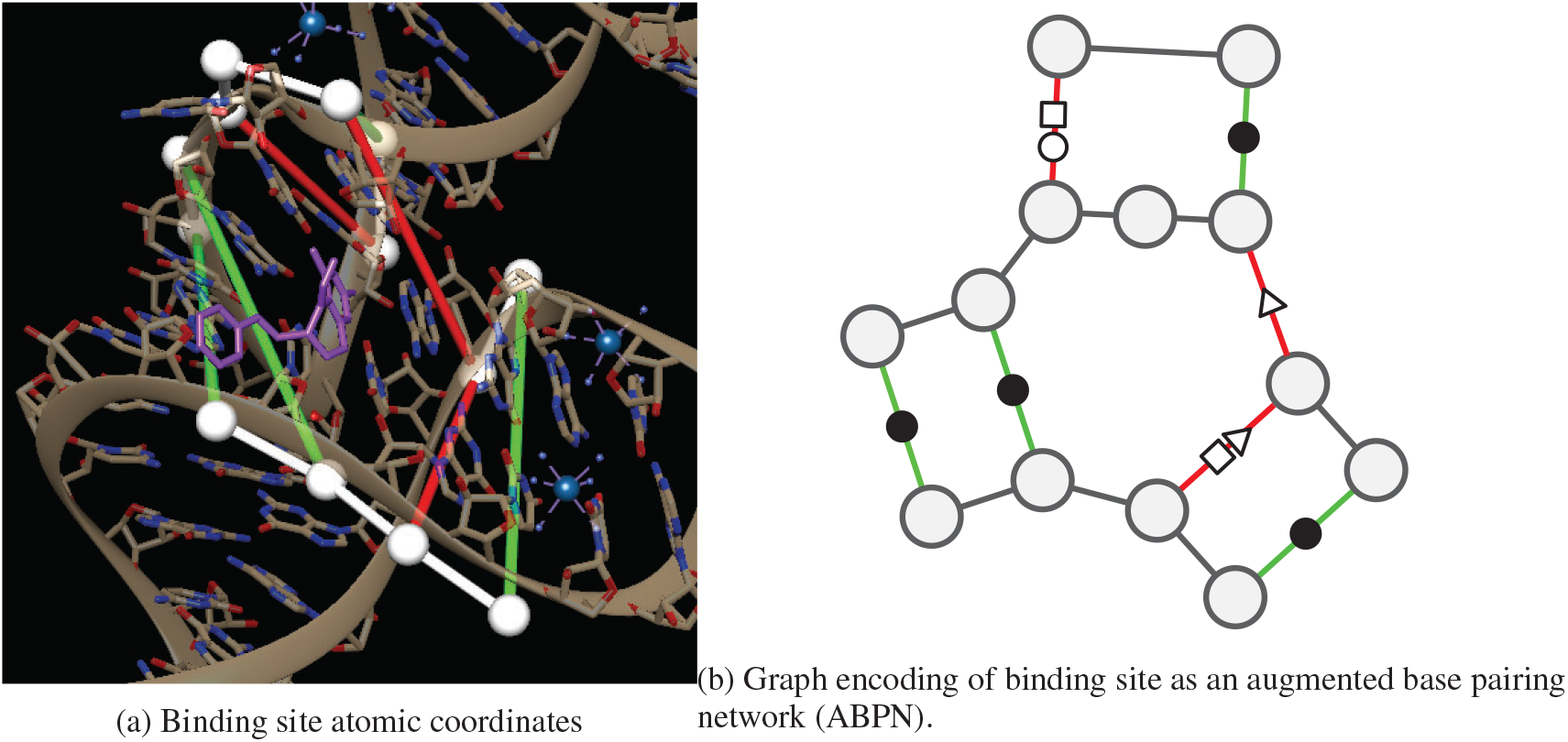
RNA structure representation of the THF riboswitch binding site (PDB: `4LVV`) as atomic coordinates using UCSF Chimera [24](left) and resulting augmented base pairing network (ABPN) (right). We superpose the ABPN in the 3D visualization. Nodes are drawn as white spheres, backbone connections are in white, and canonical and non-canonical base pairs are green and red tubes respectively. We color the edges simply to guide the eye to the corresponding base pairs but note that edge color has no special meaning to our graphs. We annotate the graph representation with the standard Leontis-Westhof nomenclature for pairing type symbols. In this case, the binding site has three canonical interactions denoted (•), and three non canonicals of types (□○, ▹, □▹).

### 1.2 Structure-based Drug Discovery and RNA Base Pairing Networks

The central aim of structure-based drug discovery is to identify compounds with high affinity to a given site or set of binding sites. A natural problem to address in this context is the prediction of binding affinity from a binding site-ligand pair. Machine learning models which solve this task can be used as alternatives to computationally expensive docking simulations to screen of ligand databases for strong binders [25]. This setting is quite feasible in the protein domain as affinities and drug screens are abundant, hence various methods have been proposed [26]. Recently, some repositories of RNA small molecule data have been made public [27] however, only a handful of binding affinities are known. Two automated affinity scoring approaches have been proposed; DrugScoreRNA, and LigandRNA [28, 29], however these rely on chemical knowledge for constructing scoring functions. While more accurate yet slower RNA docking methods are still showing limited success [23, 30]. To our knowledge, no fully data-driven methods for RNA structural drug discovery have been developed.

We therefore turn to the set of RNA-small molecule binding events captured by crystallography and made publicly available in the RCSB Protein Databank. By extracting the atomic coordinates around all organic compounds bound to RNA, we are able to obtain a set of binding site-ligand pairs for machine learning. Our model, `RNAmigos`, seeks to learn the relationship between RNA structures (represented as augmented base pairing interaction networks) and ligands (encoded as vectors of chemical features). Instead of producing a scalar affinity score for a ligand-RNA complex, we output a chemical description (fingerprint) of a ligand given a binding site. An advantage of this framing is that our model would only require as input a binding site and not an enumeration of structure-ligand pairs, as is the case in the usual affinity prediction models. Such instances of the drug discovery problem have recently been proposed in the protein domain [31, 32]. By directly yielding an active compound for each binding site, identifying ligands for large sets of sites is greatly accelerated. Given the pervasiveness of RNA transcripts, such models are likely to become especially relevant in the RNA domain.

### 1.3 Contribution

`RNAmigos` brings together domain knowledge of RNA structure, currently available crystal structure data, and graph neural networks, to show that base pairing networks can be used to automatically recover ligands for RNA structures. We propose the use of Augmented Base Pairing (ABPNs) networks which encode an enriched alphabet of base pairing interactions and demonstrate that it is a necessary component for capturing binding signatures. Molecular fingerprints predicted by `RNAmigos` serve as effective ligand search tools across diverse ligand classes and shows strong performance in two different ligand screens. Additionally, we explore the use of an unsupervised graph representation learning scheme for boosting model performance in this low-data setting. This approach holds promise for applications in other structure-function prediction tasks for RNA. Our model takes the first steps toward data-driven RNA drug discovery as well as establishes the groundwork for structure-function prediction of complex RNA structures. The core implementation of `RNAmigos` is built in Pytorch [33] and DGL [34] and is available as an open source Python 3.6 software package.

## 2 Methods

At a high level, `RNAmigos` accepts a base pairing network as input and predicts a ligand fingerprint. More formally, we learn a mapping from node and edge-attributed graphs 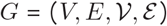 to binary vectors *ŷ* ∈ [0, 1]^*k*^ in a multi-label classification setting. The set of node attributes includes the 4 nucleotide types 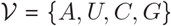 and edge attributes contain the set of 12 base pairing geometries and the backbone pair, *ε* = {`backbone`, {*cis, trans*} × {○, □, ▹}^2^}. This process is supervised by minimizing the difference between the predicted fingerprint and the observed fingerprint (in this case, co-crystallized ligand). Finally, we evaluate the quality of the predictions in a ligand screen by using *ŷ* to search for *y* in a larger set of ligands (decoys). This allows us to better contextualize the raw model performance in a more realistic setting. We provide a full overview of `RNAmigos` in **Fig. 2**.

**Figure 2:**
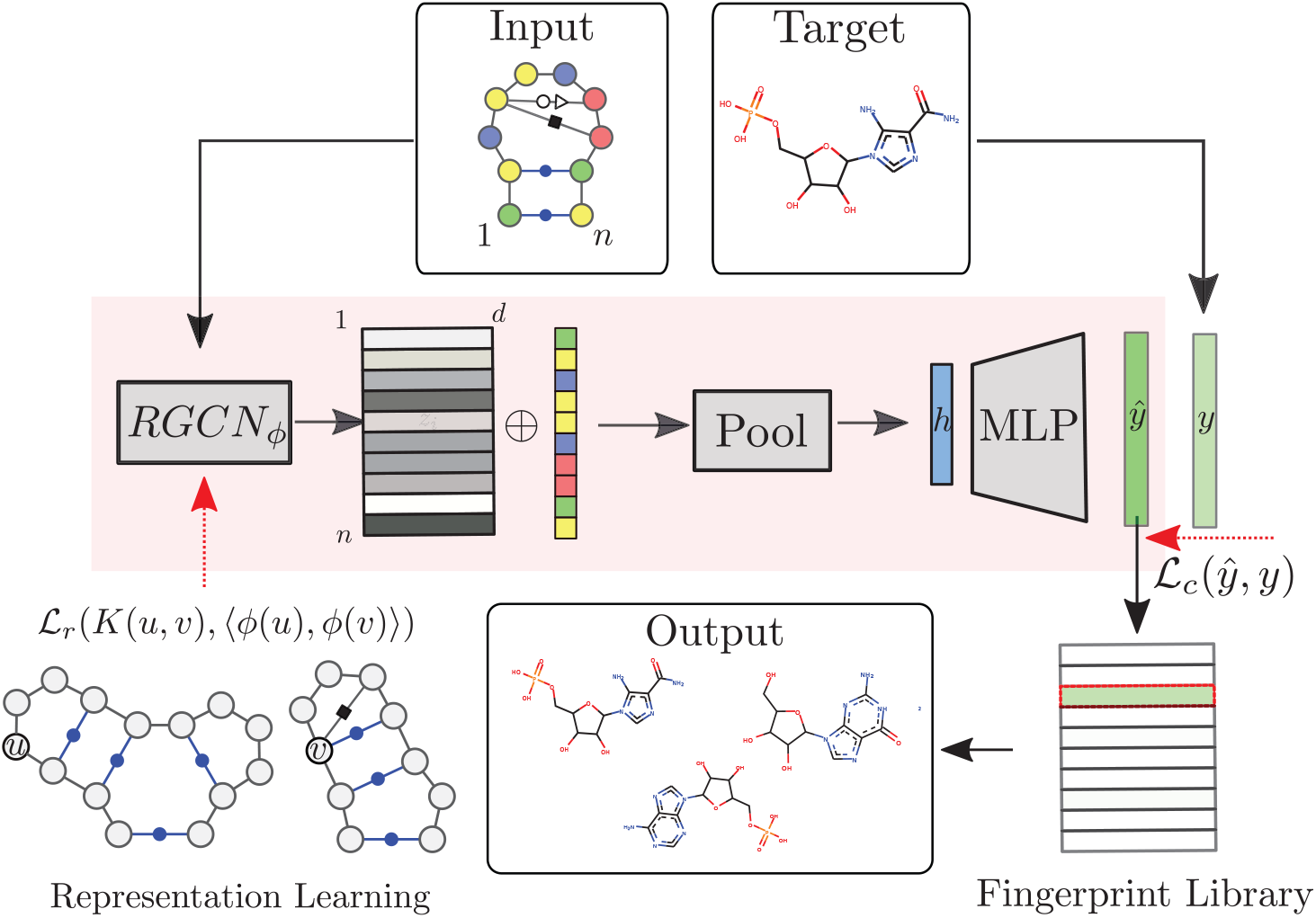
Outline of the `RNAmigos` pipeline. An base pairing network is passed as input to `RNAmigos`. In training mode, it is paired with a known ligand (Target) from which a target fingerprint *y* is constructed. The embedding network (RGCN) produces a matrix of node embeddings of dimension *n* × *d* where *n* is the number of nodes in the graph. This is followed by a pooling step which reduces node embeddings to a single graph-level vector. Finally, the graph representation is fed through a multi-layer perceptron (MLP) to produce a predicted fingerprint *ŷ*. The fingerprint is then used to search for similar ligands to the prediction in a ligand screen and thus enriches the probability of identifying an active compound. The RGCN network is pre-trained using an unsupervised node embedding framework which allows us to leverage structural patterns from a large dataset of RNA structures.

Here we outline the steps of training set construction, model architecture, and ligand screen validation.

### 2.1 Dataset Preparation

Since no benchmark datasets exist for this task, we begin by building a set of RNA-small molecule complexes from the RCSB PDB Data Bank [35]. We download all crystal structures (90% identity threshold) which contain RNA and at least one ligand. This results in 2993 PDB structures. We omit ions such as magnesium (Mg+) from the set of valid ligands as they vastly outnumber organic ligands and likely require customized models [36] We choose a maximum allowable distance between any ligand atom and any RNA atom of 10 Angstroms according to *David-Eden et al.* [18] which statistically characterized ribosome antibiotic binding sites. The number of valid sites is further reduced when we remove binding sites with fewer than five RNA residues and remove binding sites containing a large proportion of protein residues as these are unlikely to form a useful structural signal. The final training set consists of 773 binding sites associated to 270 unique ligands.

Finally, we build an ABPN from the 3D structure of each binding site identified in the previous step, where each node corresponds to a residue in the binding site, and edges are formed between edges if they form a backbone or base pair interaction. Node and edge annotations are taken from the BGSU RNA 3D Motif Atlas [16] database which maintains base pairing annotations of all PDBs with Leontis-Westhof and backbone interaction types computed by the software FR3D [37]. In this manner, each ABPN stores the nucleotide identity (`A, U, C, G`) of each of its residues a a node attribute, and each base pairing interaction corresponds to an edge with one of 13 different types (backbone + 12 base pairing geometries). Since we applied a hard distance cutoff in the crystal structure, the resulting ABPNs often have backbone discontinuities. To address this issue, we expand the ABPN to include nodes for residues found at 1-hop from the initial node set. Graphs are on average 15.76 nodes in size. At this point, the ligand is removed from the structure so that the graph contains only RNA base-pairing information. While atomic coordinates are the current source of data, we highlight that a key feature of taking ABPNs as input is that we can eventually learn from many other sources of data which are easier to obtain. A promising example comes from recent developments in predicting base pairing networks [22] from RNA sequences in high-throughput. Our model would then be able to directly take advantage of such data once it is linked to a functional label (such as a fingerprint in this case). For full details on binding site extraction and graph construction, see Supplementary Material.

### 2.2 Fingerprint Prediction

The output side of our model is a vector *ŷ* ∈ [0, 1]^*k*^ which is trained to approximate the chemical fingerprint of the bound ligand. Many functions to compute fingerprints from chemical structures have been developed, all with the common aim of encoding chemicals in a vector space where similar compounds will lie in proximal regions of the space [38, 39, 40]. In this work we use a very common fingerprint implementation known as the MACCS fingerprint [41] which has the advantage of being compact (*k* = 166 vs the usual 1024) and has interpretable dimensions. For a given chemical compound *c*, the MACCS fingerprint *f*_*c*_ is a 166 bit binary vector where each dimension indicates the presence or absence of a chemical property. For the *i*^*th*^ chemical property, *f*_*c*_[*i*] is set to 1 if the chemical property is present and is 0 otherwise. We use the set of 166 predefined chemical properties from the [42] implementation as a target vector for our model. We emphasize that the computation of the fingerprint depends only on the chemical composition of the ligand and not on the RNA binding site. In a cheminformatics setting, fingerprints are usually compared using the Tanimoto similarity, or the equivalent Jaccard distance which are simple intersections over union metrics [43] for binary vectors. However, in our machine learning setting, each fingerprint dimension is treated as multi-label category for which we output a real-valued probability. For this reason, we perform all our fingerprint comparisons using the Euclidean distance.

### 2.3 Model Architecture

Since a key feature of our ABPNs is the fact that we encode base paring geometry as an edge category (or relation type), we use a Relational Graph Convolutional Network (RGCN) [44] as the core of the fingerprint prediction model. An RGCN is a function *ϕ*_*θ*_ : *G* → ℝ^|*V*| × *d*^ with learnable parameters *θ* which maps nodes in a graph to real vectors of size *d* known as node embeddings. Embeddings for each node are obtained by repeatedly applying a trainable aggregation function over the neighbourhood of each node. The relational aspect of these networks comes in the ability to account for multi-relational graphs, (graphs with edges belonging to different categories) by allowing for parameter sharing between edge types. For this work, we consider the base pairing types to be distinct categories. We believe this is a fair approximation given the results of isostericity comparisons showing that computing the geometric discrepancy between of all pairs of edge types yields close to a diagonal matrix [45].

Once node embeddings are computed, we concatenate the resulting embedding matrix with a one-hot encoding of the input graph’s nucleotides (`A, U, C, G`). Next, graph-level representation is obtained by applying a widely used trainable Graph Attention Pooling layer [46], *GAT*_*π*_ : (*V, E*) → ℝ^*d*^ to map the node embeddings to a single vector *h* ∈ ℝ^*d*^. Finally, we feed *h* through a Multi Layer Perceptron which yields probabilities for each index of the fingerprint *y*. We supervise this process using the binary cross entropy between the predicted fingerprint and the observed over all dimensions *i*,

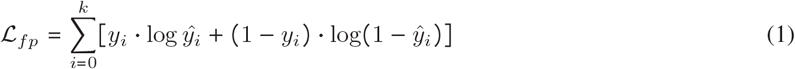

### 2.4 Unsupervised Pre-Training: ABPN Node Embeddings

Since a relatively small number of RNA-small molecule binding events have been captured by crystallography, we propose an auxilary task where data is much more abundant. More specifically, we take inspiration from recent unsupervised graph representation learning techniques [47] to leverage information from the much larger set of all crystallized RNA structures (57,6060 training points vs 773 binding sites). Instead of supervising our model using only feedback from the fingerprints, we can build an RGCN with parameters *π*′ which has learned to efficiently represent local RNA structure as vector embeddings. If this training is successful, the fingerprint prediction network can use the unsupervised RGCN weights *θ*′ as a richer initialization for the RGCN in the supervised setting (See **Fig. 2** for an illustration of this process). In the unsupervised setting, we simply require that the embeddings for a pair of nodes *ϕ*_*θ*_(*u*) and *ϕ*_*θ*_(*v*) respect a user-defined similarity relation.

We are free to choose the pairwise similarity function *K* : (*u, v*) → [0, 1] according to domain knowledge,.

We adapt the node similarity function proposed in `struc2vec` [48] which allows us to capture local structural similarity across graphs. Other node kernels such as the ones used in `GraphClust` for RNA 2D structures [49] are only able to compare nodes within the same graph and are affected by the distance between nodes which is not necessarily related to structure. Our similarity function addresses these limitations by comparing the distribution of edge types in the local neighbourhood of *u* and *v*. We define *K*_*L*_ (*u, v*) as

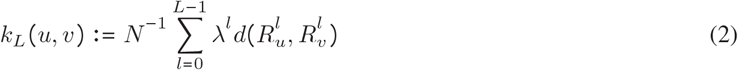

Where *d* is a function which compares the sets of edges edges at a distance *l*, from *u* and *v*, denoted 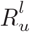, 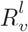. We use N as a normalization constant 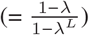 to ensure the sum saturates at 1. The *λ*^*l*^ is a decay term which allows us to attend more to structural information close to the root nodes and we set *λ* = 0.5. In our case, we define *d* to be a simple overlap measure on the histograms of base-pairing edge types *f*_*R*_, and 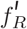 (i.e *f*_*R*_(*i*) stores the number of times edge type *i* is observed).

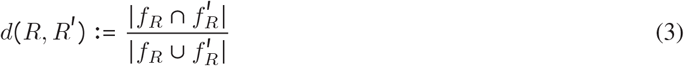

To compensate for the over-representation of a few edge types such as backbones and Watson-Crick edges, we scale the *d* value with the commonly-used Inverse-Document Frequency (IDF) factor [50].

The quality of the representations is evaluated using an L2 loss between the similarity value in the graph space and the cosine similarity of the embeddings,

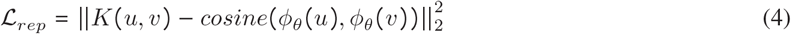

The inductive nature of graph neural networks allows us to simply input the obtained weights Φ to the different task of fingerprint prediction network. In this manner, we inject general RNA structure information in the supervised learning process where data is scarce.

### 2.5 Ligand Screen

Finally, we place the raw performance measure of the model (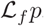, similarity to the observed ligand fingerprint) in the context of a ligand screen. In a ligand screen, one is given a set of compounds 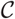 and a target. The goal is to rank ligands in 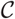 by likelihood to be active binders. Here we place our predictions in such a setting. Since no external benchmarks exist for the task of fingerprint reconstruction, this lets assess whether predictions are close enough to the true value to be useful. Given a predicted fingerprint *ŷ* ∈ [0, 1]^166^ for a binding site and a library of fingerprints 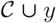, the screen ranks all 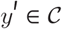 by similarity to *ŷ*. A good *ŷ* will assign a high rank to *y*, the true binder and thus is presumed to be an effective prediction for 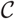. For comparison purposes, we compute a normalized rank as follows:

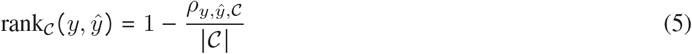

Where 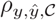 is the un-normalized rank of the true ligand *y* in 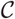 relative to the prediction *ŷ*. A successful predictor will rank the true ligand as closest to its prediction (normalized rank close to 1), while a random predictor will result in an average rank of 0.5.

Considering that the distribution of RNA ligands appears to cluster to specific sub-regions (see Supp. Fig 2), this readout also ensures that a classifier does not obtain a good score by simply predicting the average ligand as it would with a raw distance between *y* and *ŷ*.

In benchmarking settings 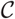 is typically known as a decoy set where the set is assumed to entirely of non-binders except for a single binder, which we hope to be able to detect. We construct two decoy sets for our experiments. Since there are currently no experimentally validated data sets of active and inactive binders for a given RNA site (such as DUDE for protein [51]), our first set consists of all RNA-binding ligands (270 ligands) in the PDB as our ligand library. The second decoy set is constructed using *DecoyFinder* [52] on default settings, which samples a list of 36 decoys for each compound such that generic chemical properties are preserved while potentially disturbing binding potential. Of course, this test assumes that chemical dissimilarity between an active compound 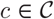 implies inactivity which is not always the case [53]. However, the current aim of our work is simply to determine whether ABPNs retain significant amount of information about its observed ligand preferences, for which this test is sufficient.

## 3 Results

We evaluate our performance using the rank metric defined in Equation 5. We report resulting rank over the list of all RNA-small molecule pairs as well as the set of all decoys for the each ligands, following the two decoy benchmark process.

Due to the limited size of our labeled data set, we performed a 10-fold cross-validation to include all training pairs in the evaluation and provide a more accurate measure of performance.

Node embeddings are computed using a 3-layer RGCN, each layer consisting of 16 dimensional inputs and outputs, a graph attention layer computing a 16 dimensional graph embedding and a fully-connected layer, which outputs a 166-dimensional vector. See Supplemental Table 1 for full model architecture and hyperparameters. Variations of the architecture used did not have strong effects on performance, so no extensive hyper-parameter search was conducted. We leave the exploration of other architecture choices for future work.

### 3.1 Augmented RNA Base Pairing Networks Encode Binding Preferences

#### Setting

The first hypothesis to test is that the proposed framework (*ABPN*) is able to retrieve some signal. To explore this question, we compute the rank and distance metrics on corrupted data. We compare this performance to three baselines:

- *random* consists of a synthetic label set where each binding site is assigned a uniformly random 166-dimensional binary vector.
- *swap* is designed to account for imbalances in the data (some ligands are more frequent than others): each binding site is assigned a fingerprint selected at random from the set of observed fingerprints. The overall distribution of ligand fingerprints thus remains the same but the input-output correlations are broken.
- *majority* is a constant ligand annotation computed as a majority vote over all fingerprints at each index. This is to be compared to the *swap* to check that the only thing that can be learnt on swapped data is over-representation of some ligands within the experiment.

The distributions of performance over each input-output pair is visualized for all experiments in **Fig. 3** as a kernel density estimate and summary statistics can be found in **Table 1** with accompanying standard deviations in Supp. Table 2. We also assessed the statistical significance of the difference of the means in a pairwise Wilcoxon rank test which is shown in **Table 2**.

**Table 1:** Mean ligand screen ranks and L2 distance achieved on held-out binding sites for each experiment and decoy set.

| Experiment | Ranks |  | L2 |  |
| --- | --- | --- | --- | --- |
|  | *DecoyFinder* | RNA | *DecoyFinder* | RNA |
| random | 0.611 | 0.542 | 0.502 | 0.502 |
| majority | 0.621 | 0.603 | 0.175 | 0.179 |
| swap | 0.617 | 0.603 | 0.177 | 0.179 |
| no-label | 0.628 | 0.606 | 0.176 | 0.180 |
| primary | 0.624 | 0.592 | 0.181 | 0.186 |
| secondary | 0.631 | 0.605 | 0.178 | 0.182 |
| ABPN | 0.695 | 0.681 | 0.155 | 0.160 |
| ABPN + unsup. | **0.735** | **0.715** | **0.145318** | **0.150189** |

**Figure 3:**
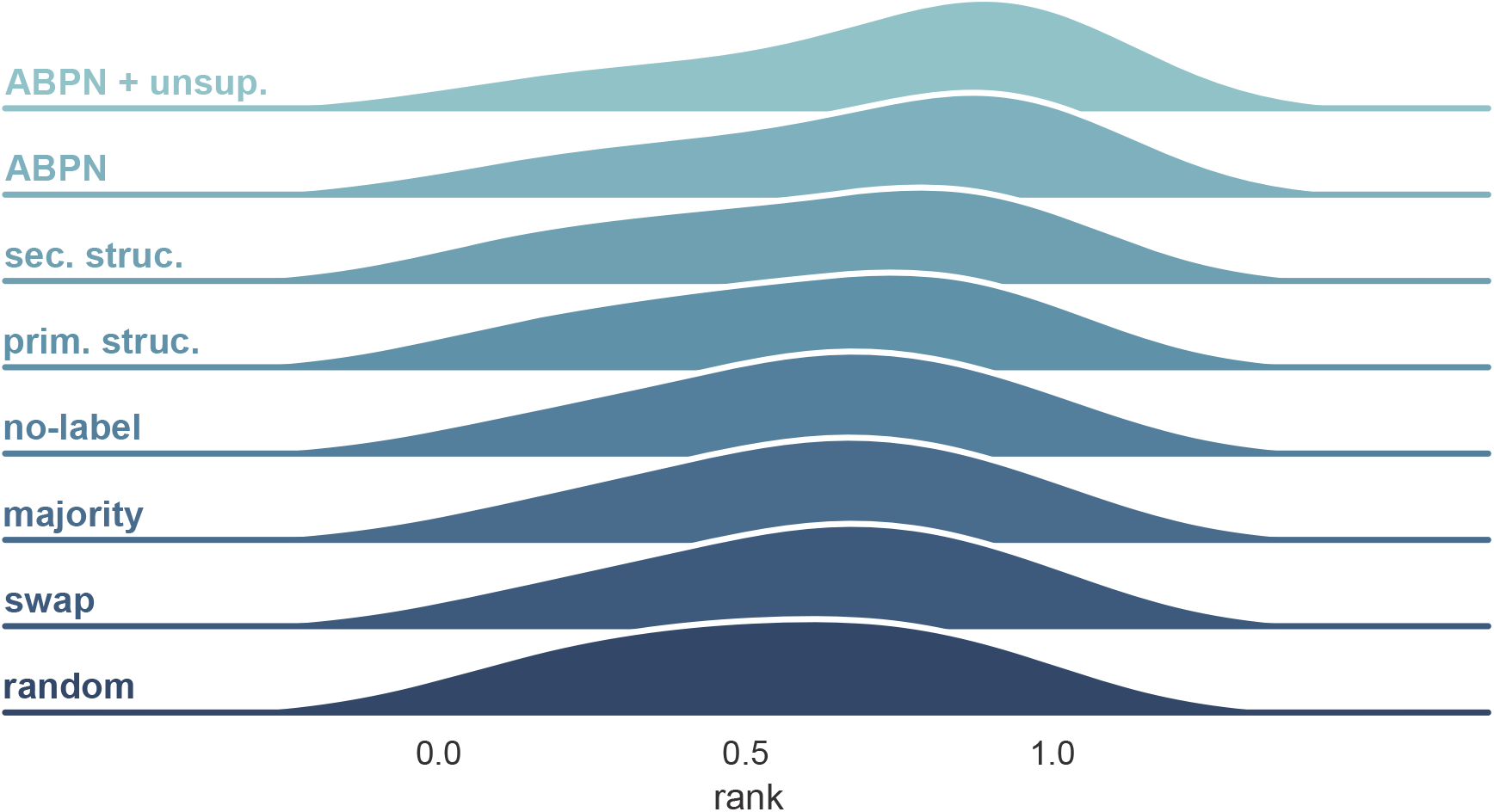
Distribution of L2 distances to the true ligand and rank achieved on ligand screening. Densities are estimated using KDE and represent the distribution achieved on all held-out binding sites during 10-fold cross-validation.

**Table 2:**
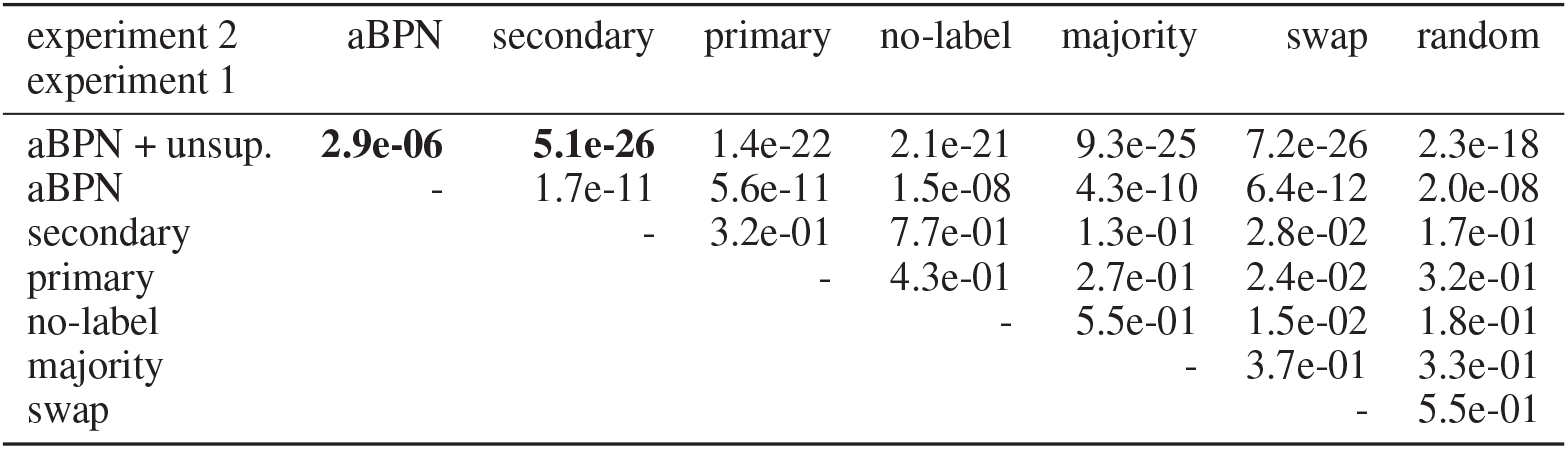
Wilcoxon rank test for all pairs of training conditions. Each entry in the table is the p-value for testing the hypothesis that the ranks resulting from a pair of experiments come from the same distribution. These are performed on the RNA decoy set. We provide the test results for the *DecoyFinder* decoy set in Supplemental Table 3 material and show consistent results.

| experiment 2<br>experiment 1 | aBPN | secondary | primary | no-label | majority | swap | random |
| --- | --- | --- | --- | --- | --- | --- | --- |
| aBPN + unsup. | **2.9e-06** | **5.1e-26** | 1.4e-22 | 2.1e-21 | 9.3e-25 | 7.2e-26 | 2.3e-18 |
| aBPN | - | 1.7e-11 | 5.6e-11 | 1.5e-08 | 4.3e-10 | 6.4e-12 | 2.0e-08 |
| secondary |  | - | 3.2e-01 | 7.7e-01 | 1.3e-01 | 2.8e-02 | 1.7e-01 |
| primary |  |  | - | 4.3e-01 | 2.7e-01 | 2.4e-02 | 3.2e-01 |
| no-label |  |  |  | - | 5.5e-01 | 1.5e-02 | 1.8e-01 |
| majority |  |  |  |  | - | 3.7e-01 | 3.3e-01 |
| swap |  |  |  |  |  | - | 5.5e-01 |

#### Performance

In the RNA-ligands setting, our full model achieves a rank of 0.68 and an mean-squared error (MSE) of 0.150 to the true fingerprint. The *random*, *swap* and *majority* experiments respectively yield ranks of 0.542, 0.603 and 0.603 and MSEs of 0.5, 0.18 and 0.18. This confirms that this model retrieves signal for the data and outperforms baselines. This conclusion is statistically significant based on a Wilcoxon p-value of at most 7*e*^−18^ between the model and the randomized results. As expected, the majority scheme is statistically equivalent to the swapped one and superior to the random one. These results are similar in the *DecoyFinder* setting where the mean rank of the model is 0.69 compared to 0.62 in the majority setting. This shows that the full model successfully retrieves some signal and beats the baselines.

### 3.2 Augmented Base Pairing Networks Encode Ligand Binding Information

We want to test the hypothesis that robust descriptors in the form of ABPN from the RNA domain knowledge are key to retrieve this signal. The question is whether the non-canonical interactions encode information that lower levels of structure (secondary, primary) do not. We answer this question by performing 3 ablation experiments on our training set:

- *primary* encodes the binding site as graphs that only contain node sequence and backbone interactions.
- *secondary* uses only information from the secondary structure which includes canonical pairs and backbones.
- *no-label* preserves all the interactions (and thus graph structure) in the graph (including non-canonical) but do not distinguish between different edge types (i.e. edges only have one label).

In all these conditions, we find that **performance is no better than the randomized baselines**, indicating that non-canonical interactions are essential for encoding specificity in ligand binding. Indeed, the best performing model is *no-label* which has a Wilcoxon p-value of 0.55 with the *majority* experiment and of 1.7*e*^−18^ with the ABPN. This finding is in agreement with biological literature on RNA binding sites and the importance of complex structural motifs for determining functional specificity [18, 7]. This is a major validation of the hypothesis that these are the correct representation for RNA structure for this task.

### 3.3 Unsupervised Pre-Training Boosts Performance

As explained before, one major limitation for this supervised task is the paucity of data. We investigated the possibility of using unsupervised learning by leveraging pre-trained weights on an unsupervised task as described above, and denote this experiment as *ABPN unsup*. The use of unsupervised pre-training of the node embedding network provides a significant performance boost over a network trained only on fingerprint reconstruction (MSE = 0.68 vs MSE=0.715), with a p-value of 2.9*e*^−6^ This is a methodological insight that can have applications for various other RNA-related tasks for which labeled data is typically scarce.

### 3.4 Our Model Can Predict Diverse Ligand Classes

Next, we ask whether the positive results can be explained by a small set of ligands, or whether it is able to achieve high scores on a diverse set of ligands. To get a better view of performance, we plot the same prediction scores but averaged over ligand types (270 unique ligands) against a hierarchical clustering dendrogram of each ligand (shown in **Fig. 4**).

**Figure 4:**
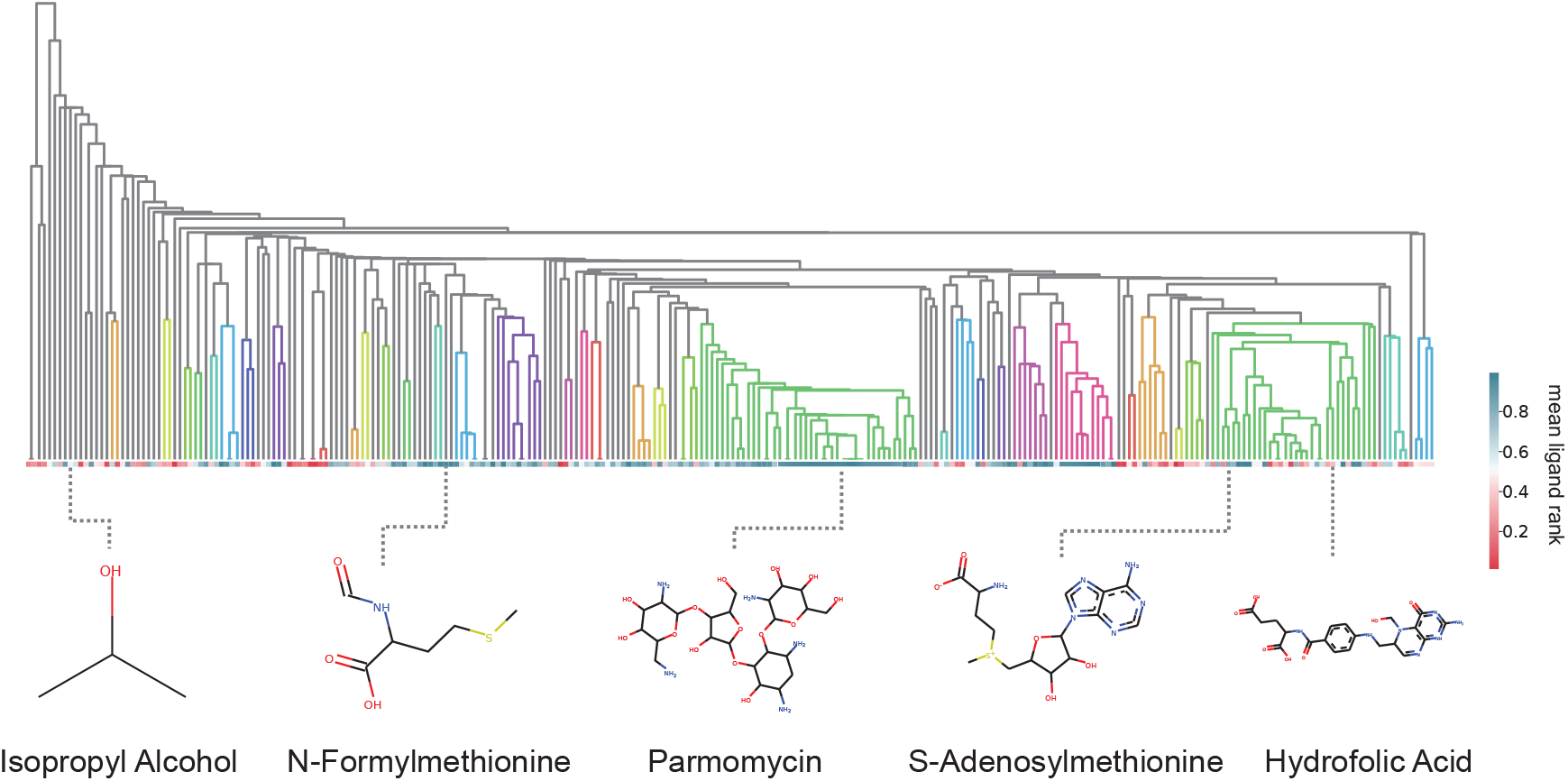
RNAmigos performance by ligand class. Hierarchical clustering dendrogram of the ligands, classifying ligand families by similarity. Each cell in the horizontal grid is the average score for binding sites containing a given ligand. Ligands belonging to the same tree are grouped together by the clustering procedure. Colored-in sub-trees denote tight clusters which contain ligands within 0.25 Jaccard distance.

Colored-in subtrees indicate groups of ligands that are similar, (i.e., within 0.25 Jaccard distance of each other) which would indicate strong clustering. In this manner, we are able to assess the performance across ‘classes’ of similar ligands. We first observe that successful classifications are not restricted to a single class of ligands and instead show good predictions for diverse ligands. Interestingly, the class that is most consistently predicted accurately corresponds to the aminoglycosides (highlighted in the pink cluster to the right). Aminoglycosides are a class of antibiotics binding to bacterial RNA with well-defined binding sites[54], and are quite abundant in the dataset. Nucleic acid-like compounds, many of which bind riboswitches, also form a large family of binders (green) however results were less consistent than for aminoglycosides. A possible explanation for strong performance on aminoglycosides, apart for the larger number of examples obtained, is that these are typically large polysaccharide-like structures with a large number of interactions with the RNA. On the other hand, riboswitches bind much smaller molecules with a limited number of interactions. As a result, binding site requirements are much more complex and specific with aminoglycosides and the large number of interactions can only be fulfilled by a limited number of molecules. Finally, ligands clustered on the left of the dendrogram show the weakest performance. Since these groups show little branching in the dendrogram, we can conclude that they represent sparsely populated ligand classes for which we have few examples and thus, obtaining more data in these regions could improve performance.

## 4 Discussion

We have developed a unique computational platform, `RNAmigos`, to show that augmented RNA base pairing networks contain useful ligand binding information. The significance of our results is two-fold.

We show for the first time that ABPNs encode sufficient information for a classification task, and establish an initial methodological primitives for such a task. To date, the majority of computational methods which leverage ABPNs have focused on sequence to structure [55, 22] prediction and motif identification [21, 16]. While these tasks involve some degree of learning, the relevance of higher-order interactions lies ultimately in their potential to specify function, which until now has been left unexplored. Interestingly, these findings come at a time when information of the type our model uses is becoming more widely available. Computational prediction tools such as [22] promise to yield large amounts of higher-order RNA pairwise interaction data without need for costly crystallography experiments. This opens the door to applying such data in other important biological problems such as RNA binding protein prediction [56] and ion binding [36]. Furthermore, the promising results obtained from the unsupervised pre-training provide a methodological building block for assisting in supervised learning on complex RNA structures.

Second, our findings take an initial step towards data-driven methods for systematically identifying drugs binding to RNA, and pinpoint ABPNs as essential for this task. The finding that only ABPNrepresentations of binding sites was able to produce a significant signal in the task indicates that richer representations are necessary for successful classification when complex interactions are at play. Since our prediction is a fingerprint vector (chemical descriptor) and not a simple classification of ligands (i.e directly selecting a single ligand as output, or predicting an affinity), the fingerprint itself can be used as a tool for searching large ligand databases. While performance was strong across different ligand classes, it is apparent that classes for which data is more abundant received more consistently positive predictions. Therefore, as more examples of RNA-ligand complexes are characterized by experimental and computational techniques, we believe that the performance of our platform will improve. Additional data will also allow for considerations regarding properties desired in medical applications such as synthesizability and drug-likeness [57]. Our choice of graphs for binding site representation reflects this consideration, as graphs can natively hold additional information such as evolutionary or chemical without requiring changes to the pipeline. Eventually, computational predictions of ABPNs from sequence [22] combined with our methods will enable transcriptome-scale searches for binding sites.

We hope that this work will motivate further investigation of the links between ABPNs and RNA function, and eventually facilitate efforts in RNA targeted drug discover.

## Acknowledgements

The authors would like to thank Mathieu Blanchette and Jacques Boitreaud for helpful feedback and discussions.

