## Supplemental Material for "Augmented base pairing networks encode RNA-small molecule binding preferences"

### 1. Data Preparation

A crucial step in our learning pipeline is the construction of ABPNs from crystal structure data. Once all crystal structures are acquired, we consider spheres of varying radii around the ligand to define a binding site. We studied two parameter choices which affect the number and quality of the extracted binding sites: radius and protein vs. RNA content. As the radius increases, we obtain a larger number of binding sites. However, since the crystal structures often contain proteins, we increase the probability that the binding site will be dominated by protein residues. We tehrefore compute the ratio of RNA to Protein residues in the binding site. The resulting counts for binding sites are shown in **Fig. ??**. From this data we choose a minimum RNA concentration of 0.6 for our training model. If a PDB contains multiple binding events of the same ligand, we keep only one at random to reduce redundancies. At this stage, we have identified a set of atomic coordinates which correspond to binding sites. Since we apply a hard distance cutoff in the crystal structure, the resulting graphs often have chain discontinuities. To address this issue, we add all 1-neighbour breadth first nodes to the original graph, as well as remove any disconnected components with fewer than 4 nodes.

### 2. Unsupervised Pre-Training

We show in **Fig. ??** and example of a pair of nodes that obtained a high similarity score after embeddings were computed.

### 3. Model Architecture and Hyperparameters

### 4. Results

| **Hyperparameter** | **Value** |
|---|---|
| RGCN Layers Dimensions | 16, 16, 16 |
| RGCN Number of Relations | 13 |
| RGCN Basis Sharing | None |
| RGCN Activation | ReLu |
| RGCN Dropout Probability | 0.5 |
| GAT Layer | Default |
| Fully Connected Dimensions | 16 166 |

Table 1. Hyperparameter choices for learning pipeline. RGCN parameters are identical for the unsupervised pre-training and the fingerprint prediction networks.

| Experiment | **Ranks** | | **L2** | |
|---|---|---|---|---|
| | *DecoyFinder* | RNA | *DecoyFinder* | RNA |
| random | 0.265880 | 0.276721 | 0.0384392 | 0.038299 |
| majority | 0.320012 | 0.269375 | 0.073969 0.074892 | |
| swap | 0.319836 | 0.269233 | 0.071212 | 0.071308 |
| no-label | 0.317259 | 0.272816 | 0.072830 | 0.073768 |
| primary | 0.323843 | 0.064917 | 0.181 | 0.066853 |
| secondary | 0.318527 | 0.299428 | 0.074738 | 0.076667 |
| ABPN | 0.322124 | 0.301635 | 0.091479 | 0.092328 |
| ABPN + unsup. | 0.303712 | 0.294309 | 0.093006 | 0.095090 |

Table 2. Standard deviation on ligand screen ranks and L2 distance achieved on held-out binding sites for each condition on both decoy sets.

| method_2<br>method_1 | ABPN | secondary | primary | no-label | majority | swap | random |
|---|---|---|---|---|---|---|---|
| ABPN + unsup - | 2.9-06 | 5.0e-26 | 1.4-22 | 2.0e-21 | 9.3e-25 | 7.1-26 | 2.3e-18 |
| ABPN | - | 1.6e-11 | 5.6e-11 | 1.4e-08 | 4.2e-10 | 6.3e-12 | 2.0e-08 |
| secondary | | - | 3.2e-01 | 7.6e-01 | 1.2e-01 | 2.8e-02 | 1.7e-01 |
| primary | | | - | 4.2e-01 | 2.7e-01 | 2.3e-02 | 3.1e-01 |
| no-label | | | | - | 5.5e-01 | 1.5e-02 | 1.7e-01 |
| majority | | | | | - | 3.6e-01 | 3.3e-01 |
| swap | | | | | | - | 5.4e-01 |

Table 3. Pairwise Wilcoxon test for the DecoyFinder decoy set over the ligand ranks.

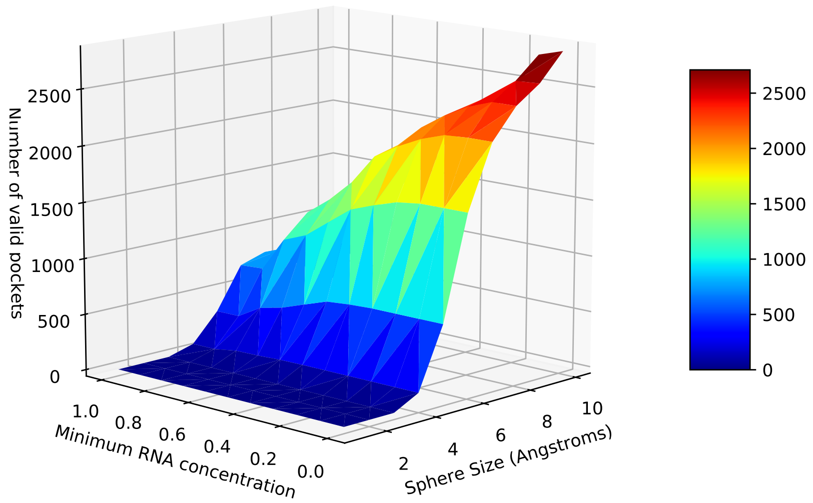


Figure 1. Number of binding sites retrieved versus distance threshold and RNA concentration threshold

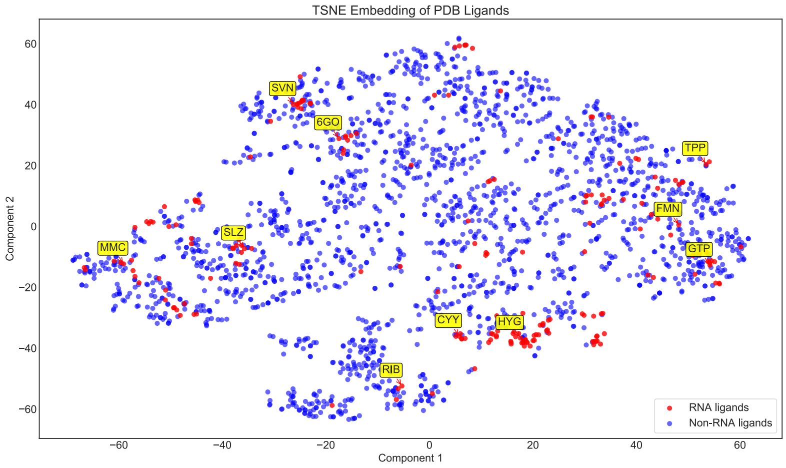


Figure 2. Two dimensional TSNE [?] embeddings of chemical fingerprints sampled from the PDB databank (RNA and protein binding). RNA ligands are highlighted in red and protein ligands in blue. We label a few interesting ligands such as 'KAN' and 'FMN' which correspond to well-known RNA binding classes known as aminoglycosides and riboswich-binding amines respectively.

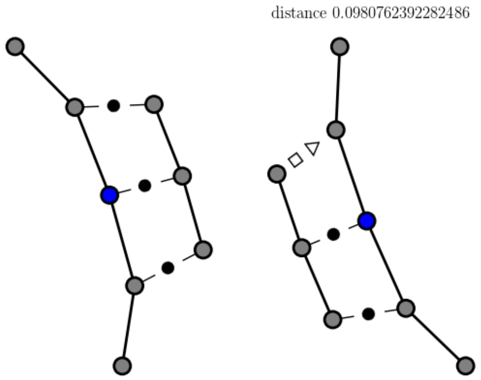


FIGURE 3. Example of pair of nodes given similar embeddings $\phi(u), \phi(v)$. The central pair of nodes which were used to make the comparison are colored in blue.
